# *Acacia gerrardii*, a desert plant as a sustainable source of natural products with antimicrobial properties

**DOI:** 10.1101/2024.09.13.612817

**Authors:** Salem Elkahoui, Ahmed Eisa Mahmoud Ghoniem, Mejdi Snoussi, Zohaier Barhoumi, Riadh Badraoui

**Author notes:** **Corresponding author. E-mail address:**, (S. Elkahoui).

## Abstract

The aim of this study was to determine the chemical composition of methanolic extracts obtained from *Acacia gerrardii* (Benth.) leaves collected from the Hail Region of Saudi Arabia, to identify new bioactive compounds, and to their antioxidant activities, antibacterial and antifungal potentials, and their mineral composition. The methanolic extract showed the highest inhibition zone against methicillin-resistant *Staphylococcus aureus* (MRSA), *Klebsiella pneumoniae* and *Escherichia coli*. This activity is dose dependent. MIC and MBC values obtained against these bacteria in the presence of the extract were 3.125 and 50 mg/mL, respectively. The antifungal activity showed low effect against all the tested fungi (MIC: 6.25 mg/mL, MBC: 25 mg/mL). GC-MS analysis revealed the presence of 15 potent compounds. 4-*O*-Methylmannose was determined as the main compound of the methanolic extract. Omega 3 and 9 fatty acids, as well as diterpene and sterols were also characterized in the extract. Forty-nine (49) compounds were identified by HR-LC/MS, the main identified classes are amino acids, alkaloids, flavonol, flavones, flavonoids, dipeptide, indoles, stilbenes, pyrans, terpenes, glycosides, and sphingolipids. Moreover, the computational study revealed that the compounds identified in *A. gerrardii* extract binds to TyrRS from *S. aureus* (1JIJ), and the secreted aspartic proteinase 1 from *C. albicans* (2QZW) receptors with acceptable affinities and interactions. The rich phytochemical composition might be the cause of the outlined pharmacokinetic properties and molecular interactions associated antimicrobial effects, as reported by both computational and *in vitro* studies.

## 1. Introduction

The exploitation of wild plants for the healing potential of their natural products dates back to ancient times. The unique healthful properties of natural products are now being recognized and intensive research has developed in recent years. Native plants in the wild and often produce higher levels of bioactive compounds in order to ward off environmental pathogens to survive harsh conditions ^1, 2^ Aromatic and medicinal plants are potential sources of antimicrobial agents due to their richness in alkaloids, anthraquinones, saponins, terpenoids, tannins and polyphenols ^3^. The Hail region in Saudi Arabia is characterized by highly diversified vegetation that are not well-studied. The majority of these plants have aromatic and medicinal potential suggested by their aroma, high essential oil and polyphenol content. Among these categories, there are cultivated and native plant species. These plants are grown for their aerial part (flowers, seeds, leaves, stems, bark) or their underground part (bulbs and roots). These Hail region plants are characterized by a wide range of biological activities such as antioxidant, antimicrobial, anti-cancer, anti-quorum sensing, anti-biofilm activities, antiseptic, analgesic, energizing, anti-inflammatory, anti-emetic antispasmodic etc. These plants represent a huge reservoir of potential beneficial compounds, attributed to secondary metabolites, that have a great diversity of chemical structures and a very wide range of biological activities. However, the evaluation of phytotherapeutic properties as antioxidant and antimicrobial remains a very interesting and useful task, especially for plants of rare or less frequent use or not known in medicine and folkloric medicinal traditions ^4^. Indeed, determination of the chemical structure of secondary metabolites is essential to understand their bioactive properties. Secondary metabolites are and remain the object of much in vivo research and in vitro research, including the search for new natural constituents such as phenolic compounds, saponosides and essential oils ^5^. In addition to their antioxidant activities, polyphenols are metal ion chelating agents (especially iron and copper cations) associated with their chemical structures. Phenolic compounds protect plants against attack by pathogens such as insects, fungi, bacteria and herbivores. Flavonoids also protect the plant against ultraviolet radiation ^6^.

*Acacia gerrardii* (Benth.) (Figure 1) is a leguminous tree and belong to the family Mimosaceae. It has a crucial role in soil protection and fertilization through symbiotic nitrogen fixation. They grow in different climatic regions and are well adapted to unfavorable ecological conditions, and therefore are an interesting genus for soil and environment protection. They are widely distributed in diverse regions through the world, especially in arid and semi-arid zones ^7^. There are 1350 species of *Acacia* in the world ^8^, distributed as follows: 144 in Africa, 185 in America, 993 in Australia and in the Pacific regions and 89 in Asia ^9^. Acacias predominate on soils containing a high proportion of sand and gravel, such as dunes, sandy plains or rocky ridges, where they form open forests scrubs ^10^ which make it an appropriate tree for Saudi soils. They are known by their ability to colonize soils with low fertility due to their ability to fix atmospheric nitrogen through their symbiotic association with Rhizobia ^10^. Due to their great biodiversity and their ability to develop a double symbiotic association with nitrogen-fixing *Rhizobium* bacteria and with endomycorrhizal fungi, acacias have gained industrial, socio-economic and ecological importance, and have attained significant potential for sustainable development ^11^.They also constitute a n important source of firewood, fiber, food, fuel, tannins, gum, handicrafts, soil stabilization, ornamentals, shelter and agroforestry system ^12, 13^. In addition, several species have been reported to be promising fodder trees, good source for production of high-quality honey, and having multiple medicinal purposes ^14–18^. Acacias also play a fundamental role in the environment protection by accelerating decarbonisation and can participate in the sustainable development of the regions where they grow.

**Figure 1.**
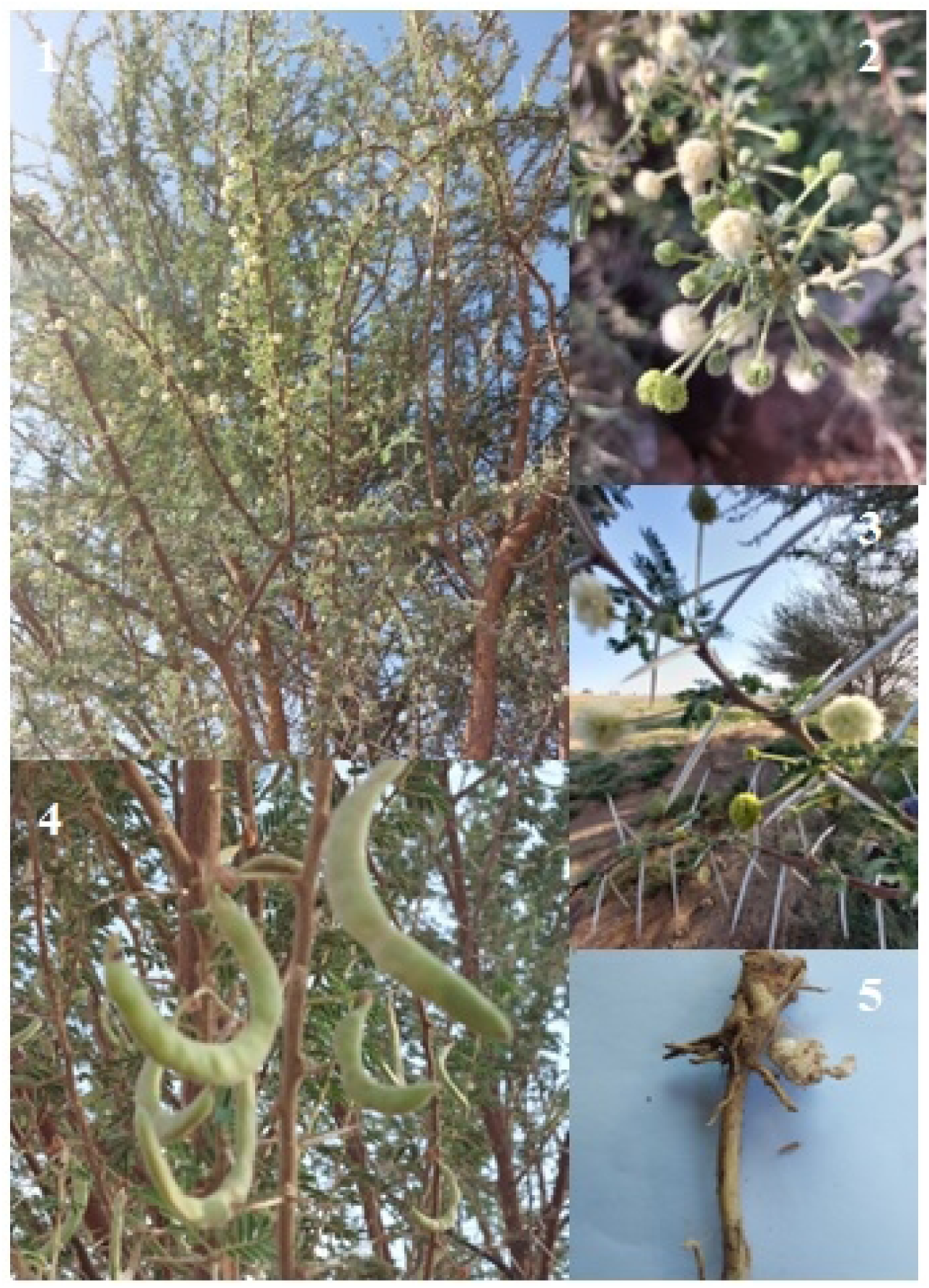
Different parts of *A. gerrardii*. 1) Overview of a tree 2) Close-up view of flowers 3) Close-up view of spines 4) Green pods 5) Nodules

The current study aims to elucidate the mineral and chemical composition, antimicrobial and antioxidant activities of methanolic extracts obtained from *A. gerrardii* leaves. The potential uses of metabolites found in the methanolic extract of *A. gerrardii* leaves for biological activity were further investigated using computational analyses, particulary through pharmacokinetic study and targeting TyrRS from S. aureus (1JIJ) and secreted aspartic proteinase 1 from *C. albicans* (2QZW) receptors.

## 2. Materials and methods

### 2.1. Material sampling

*A. gerrardii* leaves were collected in September 2022 from Hail region, Kingdom of Saudi Arabia. The specimen was recorded and deposited at the herbarium of the Department of Biology - College of Science, University of Hail, Hail - Kingdom of Saudi Arabia under a specimen voucher, AGS01. Briefly, 400 mL of methanol and 40 g of leaves were macerated for 48 hours at room temperature. The mixture was then extracted three more times using the same procedure. The extract was subjected to centrifugation at 12000g/min and the obtained supernatant was evaporated (extract yield: 10.12% ± 1.01, w/w).

### 2.2. Composition analysis

#### 2.2.1. Inductively Coupled Plasma Mass Spectrometry ICP-MS

Major and trace elements were extracted and analyzed by ICP-MS method. An operation for microwave-aided digestion was carried out in an Anton Paar Monowave 50 microwave synthesis reactor. Aliquots of 0.5 g of each sample were treated in a borosilicate glass jar (Reaction Vial G10) with 6 mL of pure HNO_3_. For the solutions examined by ICP atomic emission spectrometry, the remaining solution was diluted up to 100 g after reactors were opened to remove nitrous gases and cooled to room temperature (ICP-AES). In any case, no solid remnants were discovered, indicating that digestion was completed. Glass containers were thoroughly cleaned with nitric acid prior to the sample treatment to avoid cross-contamination. The results of the ICP-AES are provided in micrograms per gram (or parts per million) of dry weight ^19^.

#### 2.2.2. Qualitative spectrophotometric analyzes

The *A. gerrardii* methanolic extract was qualitatively tested for the presence of polyphenols, flavonoids, anthocyanin and tannins using a Cary 60 UV-Vis spectrophotometer (Agilent Technologies, Santa Clara, CA, USA) by following the protocol described by Dewanto, Wu, Adom and Liu ^20^.

#### 2.2.3. Quantative spectrophotometric analyzes GC-MS analysis

The bioactive substances in *A. gerrardii* leaves methanolic extract were identified using a Shimadzu Nexis GC-2030 Gas Chromatograph system (Kyoto, Japan) outfitted with a QP2020 NX Mass Spectrometer. In the constant flow mode, helium was used as a carrier gas at a rate of 1 mL/min. The column’s initial temperature was 70 °C. The temperature was held at this level for 2 min before being steadily raised by 10 °C to 280 °C. At an increased rate of 5 °C/min, the oven temperature was raised to 280 °C and kept there for 9 min. Helium was flowing at a rate of 1 mL/min, and the injection port temperature was 250 °C. 70 eV served as the ionization voltage. A 30-meter-long RTS volatile column achieved separation. Compounds were detected using a quadrupole mass spectrometer as they were expelled from the column. The detector had a temperature of 300 °C. Compound identification was done by analyzing the spectrum using MS data libraries WILEY8.LIB and NIST08 ^21^.

##### HR-LC/MS analysis

Agilent 324 Technologies®, USA’s UHPLC-PDA-Detector 323 Mass Spectrophotometer (HR-LCMS 1290 Infinity UHPLC System) was used to evaluate the phytochemical composition. The HiP sampler, binary gradient solvent pump, column compartment, and quadrupole time of flight mass spectrometer (MS Q-TOF) with twin Agilent Jet Stream Electrospray (AJS ES) ion sources made comprised the liquid chromatographic system. The system received 10 µL of material, separated in an SB-C18 column (2.1x50 mm, 1.8-particle size; Agilent Technologies). Acetonitrile and 1% formic acid in deionized water were utilized as solvents A and B, respectively. A 0.350 mL/min flow rate and MS Q-TOF were used for MS detection. Distinctive mass fragmentation patterns were used to identify compounds. Compound Discoverer 2.1, ChemSpider, and PubChem were used as primary tools to identify the phytochemical components of *A. gerradi* methanolic extract ^22^. A qualitative phytochemical investigation of the prepared Methanolic extract of *A. gerrardii* leaves was conducted using conventional methods. The outcomes were qualitatively expressed by using either positive (+) or negative ionization mode.

### 2.3. Antimicrobial activity

#### 2.3.1. Determination of the growth inhibition zone

The methanolic extract from A. gerrardii collected from Hail region was tested for its antibacterial and antifungal activities by using both disk diffusion and microdilution assays ^21, 23^.

To calculate the diameter of growth inhibition zone, pure colonies from bacterial culture cultivated on Mueller-Hinton agar medium were used to prepare a suspension of *Escherichia coli*, *Enterobacter faecalis*, *Staphylococcus hominis*, *S. aureus*, *S. epidermidis*, *Klebsiella pneumoniae*, *Pseudomonas aeruginosa*, *Acinetobacter baumannii*, and Methicillin-resistant *S. aureus*. Four yeast strains were also tested on Sabouraud dextrose agar plates (Candida utilis ATCC 9255, C. guillermondii ATCC 6260, C. tropicalis ATCC 1362, and C. albicans ATCC 20402). A stock solution from A. gerrardii methanolic extract was prepared at 100 mg/mL in DMSO-5% solution ^24^. Agar plates were inoculated with bacterial and fungal strains using a cotton swab technique. The extract was used to impregnate sterile Whatman filter paper at different concentrations (1 mg/disc, 2 mg/disc, and 3 mg/disc). The experiment was done in triplicate and the mean diameter of the inhibition zone was calculated (mGIZ) was calculated and expressed in mm. Ten microliters from a stock solution (10 mg/mL) of ampicillin and amphotericin B were used as control. The scheme proposed by Parveen and colleagues ^25^ was used to interpret the recorded results.

#### 2.3.2. Determination of MIC/MBC and MFC values

To determine the minimal concentration to inhibit bacterial and fungal growth and the minimal concentration to kill them, we used the microdilution assay on 96 well plates. For the experiment, we prepared a mother solution at 200 mg/mL in DMSO-5%. This solution was serially diluted and used to inoculate the 96 well-microtiter plates. Each well contains 95 µL of broth media (Lauria Bertani for bacteria, and Sabouraud broth for Candida strains), 100 µL from the diluted extract, and 5 µL from the bacterial/fungal suspension. The final volume in each well was about 200 µL. After incubation at 37 °C for 24 h, minimal inhibitory concentration (MIC) was defined as the minimum concentration responsible for no visible growth in the well. The concentration at which the bacteria/Candida was completely killed is defined as the minimum bactericidal/fungicidal concentration. MIC/MBC and MFC/MIC ratios were calculated, and results were interpreted to determine the nature of the tested extract using the scheme proposed by Gatsing et al. (2009) ^26^ and La et al. (2008) ^27^.

### 2.4. Antioxidant activity

#### 2.4.1. Ferric reducing antioxidant power assay (FRAP)

The reducing power was evaluated based on the reported method ^28^. For this, 100, 200, 500, 750, and 1000 µg/mL of *A. gerrardii* extract were diluted with 2.5 mL of sodium phosphate buffer (pH 6.6) and 2.5 mL of potassium ferricyanide at 200 mmol/L and 1%, respectively. After shaking, the mixture was incubated at 50 °C for 20 min. Then, 2.5 mL of trichloroacetic acid (10%) was added to the solution. The mixture vortexed for 20 seconds, then centrifuged for 8 min at 1000 rpm. Finally, distilled water (2.5 mL) and 1% ferric chloride (0.5 mL) was added to the vessel. The absorbance of each sample was measured spectrophotometrically at 700 nm and IC_50_ value of the extract was determined.

#### 2.4.2. DPPH assay

The total radical scavenging capacity of *A. gerrardii* methanolic extract was determined by using 2, 2-diphenyl-1-picrylhydrazyl (DPPH) method ^29^. The DPPH solution was mixed in the range of 1, 10, 100, and 200 g/mL of the methanolic extract. Following the recording of DO values at 515 nm, spectrophotometrically, IC_50_ was determined from the graph.

### 2.5. In silico analyzes

#### 2.5.1. Computational assay and interactions analyses

The identified phytochemicals of *A. gerrardii* were processed for the computational approach to decipher the molecular interactions with some key macromolecules towards determing their antibacterial and antiviral potentials. The 3D structures of the *A. gerrardii* compounds were retrieved from the Pubchem website or drawn using ChemDraw Pro 12.0 software package. The 3D crystal structure of TyrRS from *S. aureus* (pdb ID: 1JIJ) and the secreted aspartic proteinase 1 from *C. albicans* (pdb ID: 2QZW) receptors were collected from the RCSB PDB. Both ligands and receptors were prepared before being minimized ^30–32^. The complex units (ligands and receptors) were subjected to force field of CHARMm’s type as previously described following the selection of some key residues within as part of the grid box by ^30, 33, 34^.

#### 2.5.2. Bioavailability and pharmacokinetics

Pharmacokinetics and bioavailability parameters of *A. gerrardii* identified phytochemicals have been explored by computational analyses as previously described ^34, 35^. The analytic assessment was based on the ADMET (for absorption, distribution, metabolism, excretion and toxicity) measurements ^33^.

### 2.6. Statistical analysis

All measurements were done in triplicate and the results were presented as mean values ± SD (standard deviations) ^36^. Duncan’s multiple-range tests for means with a 95% confidence interval (*p* ≤ 0.05) was used to calculate the differences in means

## 3. Results and discussion

### 3.1. Mineral composition

The age, species and varieties of the plant as well as soil traits, climatic or seasonal condition influence the mineral composition of plants ^37^.

The concentrations of 20 different elements (Ag, Al, As, Ba, Be, Cd, Co, Cr, Cu, Fe, Mn, Mo, Ni, Pb, Sb, Se, Sr, Ti, V, Zn) were determined using the ICP-AES technique. Mineralogical analysis showed the presence of iron as the main element with 6.67 mg/g, followed by aluminum (4.70 mg/g), strontium (62.4 µg/g), manganese (34.94 µg/g), copper (20.82 µg/g), zinc (13.18 µg/g), and silver (11.24 µg/g). The presence of chromium, molybdenum, nickel, lead, antimony, selenium, titanium, and vanadium with a concentration lower than 1.58 µg/g. On the other hand, 5 elements (arsenic, barium, beryllium, cadmium, and cobalt) were not detected in the leaves methanolic extract.

The results obtained from the study by <u>Alatar, El-Sheikh, Thomas, Hegazy and El</u> Adawy ^38^ support the view that the mineral composition of the plant can change under the influence of environmental conditions. Additionally, the elemental content of *Acacia* species infected by mistletoe in Saudi Arabia has been reported to be significantly reduced in some species but A. Gerrardii was less affected compared to uninfected trees, especially in terms of potassium and sodium ^39^.

### 3.2. Chemical composition

In the current work, the phytochemical characterization of the methanolic extract of *A. gerrardii* leaves was first investigated by qualitative spectrophotometric analysis, followed by detailed quantitative chromatographic analyzes to confirm the obtained data. It was determined that the amount of phenolic compounds in *A. gerrardii* was found to be more abundant than other phytochemical groups (316.61 ± 1.21 mg GAE/g extract). Phenolic compounds were followed by anthocyanins (97.23 ± 1.10 mg cyanin chloride/g extract), tannins (19.25 ± 1.03 mg TAE/g extract) and flavonoids (11.17 ± 1.76 mg QE/g extract), respectively.

The methanolic extract from the *A. gerrardii* leaves was screened for the detection of different classes of aromatic biomolecules by GC-MS (Table 1, Figure 2). The chromatograms exhibited the peaks of the identified compounds with their respective chemical structures.

**Figure 2.**
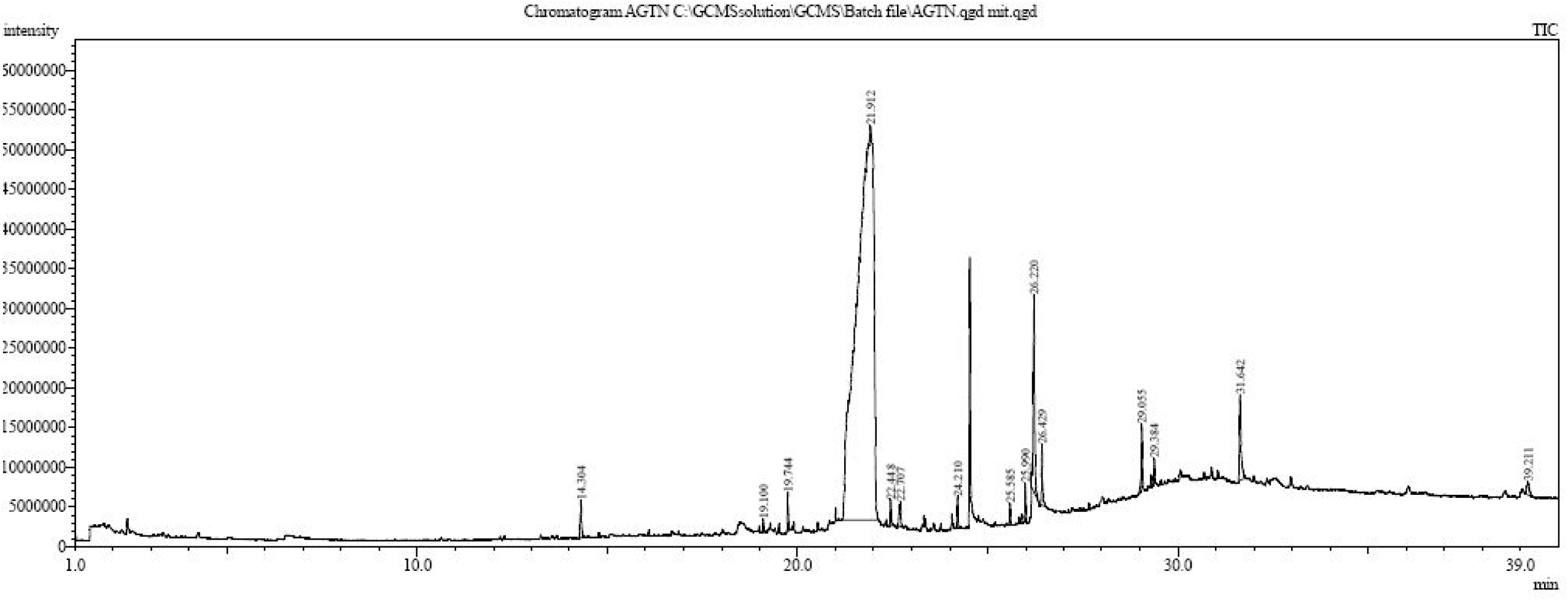
Chromatogram of the methanolic extract of *A. gerrardii* obtained through GC/MS analysis

**Table 1.** Phytochemical composition of *A. gerrardii* leaves methanolic extract using GC-MS technique (H: Hail)

| No | Compound | Class | Rt (min) | Area (%) | Formula | MassPeaks |
| --- | --- | --- | --- | --- | --- | --- |
| 1 | Azulene | Polycyclic aromatic hydrocarbons | 14.3 | 0.61 | C<sub>10</sub>H<sub>8</sub> | 257 |
| 2 | 1-(4-Ethoxyphenyl) propan-1-ol | NI | 19.1 | 0.14 | C<sub>11</sub>H<sub>16</sub>O<sub>2</sub> | 262 |
| 3 | Benzoic acid, 4-ethoxy-, ethyl ester | Carboxylic acids/Benzoates | 19.74 | 0.45 | C<sub>11</sub>H<sub>14</sub>O<sub>3</sub> | 232 |
| 4 | 4-*O*-Methylmannose | Carbohydrates/Methylmannosides | 21.91 | 73.83 | C<sub>7</sub>H<sub>14</sub>O<sub>6</sub> | 295 |
| 5 | Tetradecanoic acid | Fatty acids/Myristic acids | 22.44 | 0.3 | C<sub>14</sub>H<sub>28</sub>O<sub>2</sub> | 279 |
| 6 | Octadecamethylcyclononasiloxane | NI | 22.7 | 0.48 | C<sub>18</sub>H<sub>54</sub>O<sub>9</sub>Si<sub>9</sub> | 348 |
| 7 | *n*-Hexadecanoic acid (Palmitic Acid) | Fatty acid | 24.21 | 0.34 | C<sub>17</sub>H<sub>34</sub>O<sub>2</sub> | 337 |
| 8 | Hexadecamethylcyclooctasiloxane | Macrocyclic organosiloxane | 25.58 | 0.21 | - | 311 |
| 9 | Phytol | Acyclic diterpene | 25.99 | 0.19 | C<sub>20</sub>H<sub>40</sub>O | 269 |
| 10 | 9,12,15-Octadecatrienoic acid, (Z,Z,Z)-(Linolenic Acid) | Fatty acids, Omega-3 | 26.22 | 2.91 | C<sub>18</sub>H<sub>30</sub>O<sub>2</sub> | 358 |
| 11 | Octadecanoic acid | Fatty acids, Stearic acids | 26.42 | 0.72 | C<sub>18</sub>H<sub>36</sub>O<sub>2</sub> | 298 |
| 12 | Tetracosamethyl-cyclododecasiloxane | NI | 29.05 | 2.37 | C<sub>24</sub>H<sub>72</sub>O<sub>12</sub>Si<sub>12</sub> | 295 |
| 13 | Hexadecanoic acid | NI | 29.38 | 0.13 | C<sub>16</sub>H<sub>32</sub>O<sub>2</sub> | 296 |
| 14 | 13-Docosenamide, (Z)- | Fatty acids, Erucic acids/Omega-9 | 31.64 | 1.61 | C<sub>22</sub>H<sub>43</sub>NO | 341 |
| 15 | Stigmasterol | Lipids, Sterol | 39.21 | 0.33 | C<sub>29</sub>H<sub>48</sub>O | 308 |

NI: Non identified

The data in Table 1 shows 15 compounds by GC-MS, including 4-*O*-methylmannose (73.83%), linolenic acid (2.91%), tetracosamethyl-cyclododecasiloxane (2.37%), and 13-Docosenamide, *(Z)* (1.61%).

In addition to the volatile phytochemicals presented in Table 1, the non-volatile compounds were determined by LC-MS in the *A. gerrardii* leaf extract (Table 2, Figure 3). As evident from the data presented in Table 2, LC-MS analysis of the *A. gerrardii* leaf extract resulted in the presence of secondary metabolites such as alkaloids, carboxylic acids, flavonoids, terpenoids, in addition to well-known primary metabolites.

**Figure 3.**
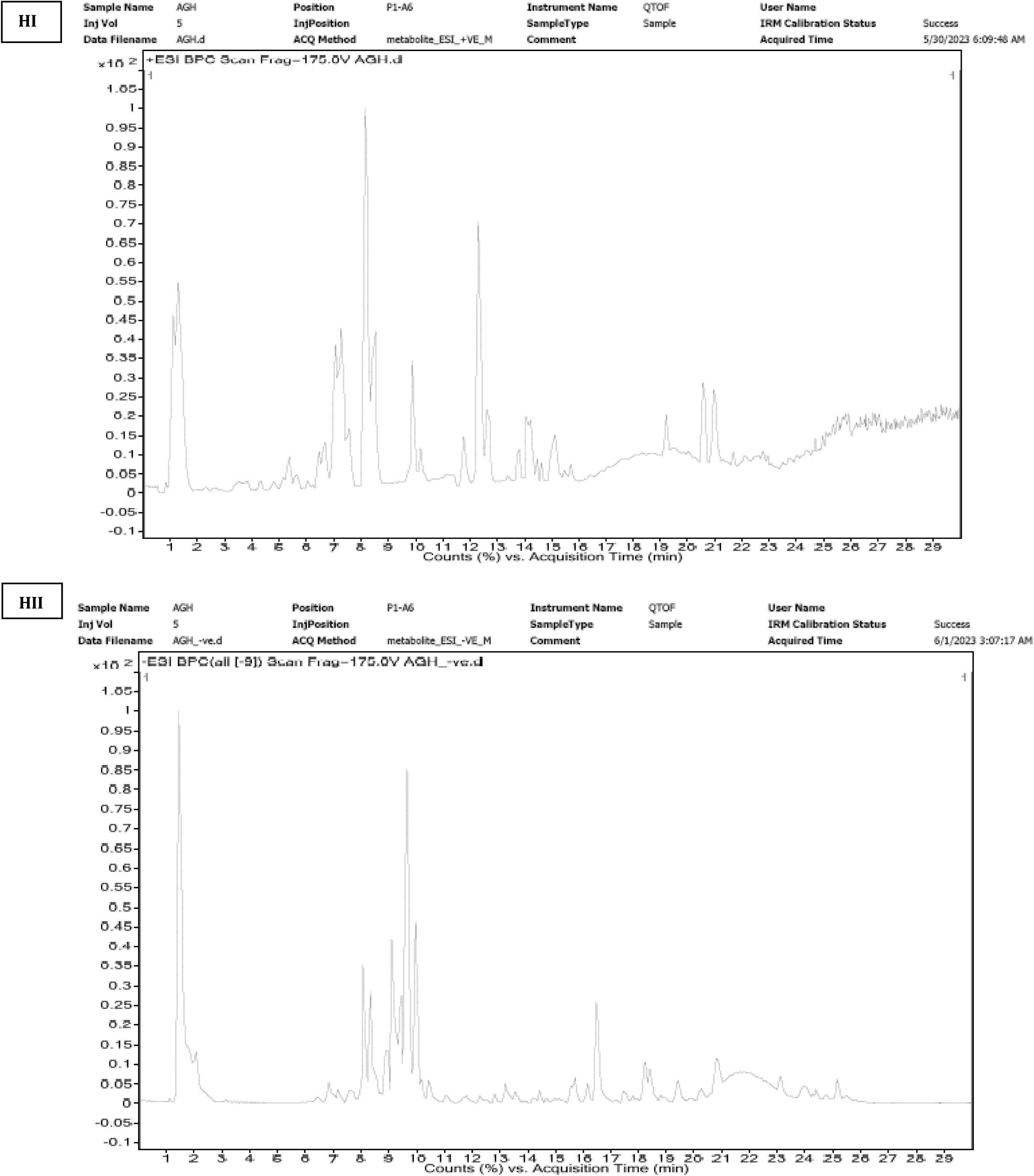
Chromatogram of the methanolic extract of *A. gerrardii* obtained through HR- LCMS analysis (H: Hail, HI: positive mode, HII: negative mode)

**Table 2.** Phytochemical composition of *A. gerradii* leaves methanolic extract using HR-LCMS.

| No | Compound | Class | Rt (min) | Mass | Formula | [M + H]<sup>-</sup> (m/z) | [M + H]<sup>+</sup> (m/z) |
| --- | --- | --- | --- | --- | --- | --- | --- |
| 1 | *L-trans*-5-Hydroxy-2-piperidinecarboxylic acid | Amino Acid | 1.044 | 145.073 | C<sub>6</sub> H<sub>11</sub> N O<sub>3</sub> |  | 146.0803 |
| 2 | Retronecine | Pyrrolizidine Alkaloids | 1.125 | 155.0932 | C<sub>8</sub> H<sub>13</sub> N O<sub>2</sub> |  | 156.1005 |
| 3 | (x)-1-Nonen-3-yl acetate | Carboxylic ester | 5.312 | 184.1473 | C<sub>11</sub> H<sub>20</sub> O<sub>2</sub> |  | 207.1366 |
| 4 | Propyl levulinate | Carboxylic acid | 5.463 | 158.0953 | C<sub>8</sub> H<sub>14</sub> O<sub>3</sub> |  | 181.0845 |
| 5 | 2-Carboxy-4-dodecanolide | Gamma-lactone | 5.694 | 242.1513 | C<sub>13</sub> H<sub>22</sub> O<sub>4</sub> |  | 265.1414 |
| 6 | Quercetin | Flavonol | 6.323 | 302.0404 | C<sub>15</sub> H<sub>10</sub> O<sub>7</sub> |  | 303.0477 |
| 7 | Phaseolic acid | Hydroxycinnamic acid | 6.329 | 296.053 | C<sub>13</sub> H<sub>12</sub> O<sub>8</sub> |  | 319.0422 |
| 8 | Cyclic dehydropoxanthine futasine | Aromatic carboxylic acid | 6.422 | 294.0739 | C<sub>14</sub> H<sub>14</sub> O<sub>7</sub> |  | 317.0632 |
| 9 | 7-(5-*O*-phosphono-α-D-ribofuranosyl)-7H-purine | Nucleotide | 6.573 | 332.051 | C<sub>10</sub> H<sub>13</sub> N<sub>4</sub> O<sub>7</sub> P | | 333.0582 |
| 10 | Cinnamodial | Aldehydes tertiary alcohol | 6.577 | 308.1622 | C<sub>17</sub> H<sub>24</sub> O<sub>5</sub> |  | 331.1514 |
| 11 | Tyrosyl-Serine | Dipeptide | 6.596 | 268.1062 | C<sub>12</sub> H<sub>16</sub> N<sub>2</sub> O<sub>5</sub> |  | 291.0953 |
| 12 | Mahaleboside | Glycoside and a member of coumarins | 7.162 | 324.0843 | C<sub>15</sub> H<sub>16</sub> O<sub>8</sub> |  | 347.0736 |
| 13 | Caulerpin | Indoles | 7.912 | 398.1262 | C<sub>24</sub> H<sub>18</sub> N<sub>2</sub> O<sub>4</sub> | 457.1401 |  |
| 14 | DL-3,4-Dihydroxymandelic acid | Acids, Carbocyclic Mandelic Acids | 8.108 | 184.0386 | C<sub>8</sub> H<sub>8</sub> O<sub>5</sub> | 183.0314 |  |
| 15 | Gnemonol A | Stilbenes | 8.61 | 696.1983 | C<sub>42</sub> H<sub>32</sub> O<sub>10</sub> | 755.2122 |  |
| 16 | Glucobrassicin | Glycosides/Glucosinolates | 8.614 | 448.061 | C<sub>16</sub> H<sub>20</sub> N<sub>2</sub> O<sub>9</sub> S<sub>2</sub> | 493.0592 |  |
| 17 | Pedaliin | Flavones and Flavonols | 8.669 | 478.1161 | C<sub>22</sub> H<sub>22</sub> O<sub>12</sub> | 477.1089 |  |
| 18 | Copalliferol B | Stilbenoids | 8.813 | 680.2027 | C<sub>42</sub> H<sub>32</sub> O<sub>9</sub> | 739.2168 |  |
| 19 | Isorhamnetin 3-rutinoside 4'-rhamnoside | Flavonoids | 8.863 | 770.235 | C<sub>34</sub> H<sub>42</sub> O<sub>20</sub> | 769.2281 |  |
| 20 | 11-*O*-Galloylbergenin | Pyrans | 8.906 | 480.0955 | C<sub>21</sub> H<sub>20</sub> O<sub>13</sub> | 479.0881 |  |
| 21 | Chartreusin | Glycoside and a benzochromenone | 8.922 | 640.1719 | C<sub>32</sub> H<sub>32</sub> O<sub>14</sub> | 639.165 |  |
| 22 | Allivcin | Flavonoids and a glycoside | 8.97 | 610.1599 | C<sub>27</sub> H<sub>30</sub> O<sub>16</sub> | 609.1528 |  |
| 23 | Rhamnetin 3-laminaribioside | Flavonoids | 9.248 | 640.1709 | C<sub>28</sub> H<sub>32</sub> O<sub>17</sub> | 639.1635 |  |
| 24 | 10-Acetoxyoleuropein | Terpene glycoside | 9.287 | 598.1964 | C<sub>27</sub> H<sub>34</sub> O<sub>15</sub> | 597.1893 |  |
| 25 | Nicotiflorin | Flavones and Flavonols | 9.316 | 594.1649 | C<sub>27</sub> H<sub>30</sub> O<sub>15</sub> | 593.1576 |  |
| 26 | Isorhamnetin 3-O-[b-D glucopyranosyl-(1->2)-a-L rhamnopyranoside] | Carbohydrate derivative<br>Glycosyl compound | 9.376 | 624.1762 | C<sub>28</sub> H<sub>32</sub> O<sub>16</sub> | 623.1689 |  |
| 27 | CMP-N-acetylneuraminic acid | Amino sugars/Neuraminic acids | 9.39 | 614.1451 | C<sub>20</sub> H<sub>31</sub> N<sub>4</sub> O<sub>16</sub> P | 659.1452 |  |
| 28 | Catechin-(4alpha->8)-gallo catechin-(4alpha->8)-catechin | Flavonoid | 9.409 | 898.1903 | C<sub>45</sub> H<sub>38</sub> O<sub>19</sub> | 957.2045 |  |
| 29 | Myricitrin | Glycosyloxyflavone (Flavonoids) | 9.414 | 464.1009 | C<sub>21</sub> H<sub>20</sub> O<sub>12</sub> | 463.0936 |  |
| 30 | (L-Seryl)adenylate | Amino acids | 9.416 | 434.0906 | C<sub>13</sub> H<sub>19</sub> N<sub>6</sub> O<sub>9</sub> P | 493.1043 |  |
| 31 | CMP-N-glycolylneuraminate | Glycosylamines | 9.666 | 630.1403 | C<sub>20</sub> H<sub>31</sub> N<sub>4</sub> O<sub>17</sub> P | 689.1563 |  |
| 32 | Rhamnazin 3-sophoroside | Flavonoids | 9.679 | 654.1868 | C<sub>29</sub> H<sub>34</sub> O<sub>17</sub> | 653.1797 |  |
| 33 | Temocaprilat | Alpha-amino acid | 9.902 | 448.1063 | C<sub>21</sub> H<sub>24</sub> N<sub>2</sub> O<sub>5</sub> S<sub>2</sub> | 447.097 |  |
| 34 | Precarthamin | Glycosides | 9.929 | 956.2321 | C<sub>44</sub> H<sub>44</sub> O<sub>24</sub> | 955.2242 |  |
| 35 | 4',8-Dimethylgossypetin 3-glucoside | Flavonoids | 9.943 | 508.1273 | C<sub>23</sub> H<sub>24</sub> O<sub>13</sub> | 507.12 |  |
| 36 | Tenitramine | Amines | 10.226 | 416.1157 | C<sub>10</sub> H<sub>20</sub> N<sub>6</sub> O<sub>12</sub> | 461.1144 |  |
| 37 | Methyl 4,6-di-O-galloyl-beta-Dglucopyranoside | Triterpenoid saponin | 10.238 | 498.0963 | C<sub>21</sub> H<sub>22</sub> O<sub>14</sub> | 497.0911 |  |
| 38 | Canescein | Cardenolide glycoside | 10.335 | 566.2647 | C<sub>29</sub> H<sub>42</sub> O<sub>11</sub> | 625.2788 |  |
| 39 | C16 Sphinganine | Sphingolipids | 12.307 | 273.2647 | C<sub>16</sub> H<sub>35</sub> N O<sub>2</sub> |  | 274.2721 |
| 40 | Juvenile hormone III | Isoprenoids (sesquiterpenes) | 12.477 | 266.1878 | C<sub>16</sub> H<sub>26</sub> O<sub>3</sub> |  | 289.177 |
| 41 | Palmitic amide | Fatty Acyls/Primary amides<br>(Fatty Acids/Palmitic Acids) | 14.183 | 255.2561 | C<sub>16</sub> H<sub>33</sub> N O |  | 278.2455 |
| 42 | 5-Acetoxydihydrotheaespirane | Oxolanes | 14.501 | 254.1885 | C<sub>15</sub> H<sub>26</sub> O<sub>3</sub> |  | 277.1777 |
| 43 | 18-Nor-4(19),8,11,13-abietatetraene | Isoprenoid/Terpenoid | 14.933 | 254.2064 | C<sub>19</sub> H<sub>26</sub> |  | 277.1956 |
| 44 | Geranyl 2-ethylbutyrate | Fatty alcohol esters | 15.091 | 252.2091 | C<sub>16</sub> H<sub>28</sub> O<sub>2</sub> |  | 275.1983 |
| 45 | Irinotecan | Alkaloids | 19.858 | 586.2779 | C<sub>33</sub> H<sub>38</sub> N<sub>4</sub> O<sub>6</sub> |  | 609.2672 |
| 46 | Oleamide | Fatty amides/ Primary amides | 20.475 | 281.2696 | C<sub>18</sub> H<sub>35</sub> N O |  | 282.277 |
| 47 | Geranylfarnesyl diphosphate | Isoprenoids (sesterterpenes) | 20.885 | 518.2615 | C<sub>25</sub> H<sub>44</sub> O<sub>7</sub> P<sub>2</sub> | 577.2756 |  |
| 48 | Ganoderic acid F | Fatty acids/Heptanoic acid | 20.872 | 570.2834 | C<sub>32</sub> H<sub>42</sub> O<sub>9</sub> |  | 593.2728 |
| 49 | Phosphatidylglycerol | Glycerophosphoglycerol | 22.25 | 818.5133 | C<sub>46</sub> H<sub>75</sub> O<sub>10</sub> P | 863.5119 |  |

As far as our literature survey could ascertain, no study has been found in the literature investigating the chemical composition of *A. gerrardii*. Therefore, it is thought that the data presented in this section of the current study will fill an important gap in the literature. However, there are some studies in the literature investigating the phytochemical compositions of other *Acacia* species. As understood from these studies, *Acacia* species stand out with the presence of phentolamine ^40, 41^, steroids ^42, 43^, tannins, alkaloids, anthraquinones ^42^, polyphenols ^42, 44, 45^, saponins ^42, 46, 47^, flavonoids ^42, 48–56^, phenolic acids ^51^, and terpenoids ^57^. <u>Batiha, Akhtar, Alsayegh, Abusudah, Almohmadi,</u> Shaheen, Singh and De Waard ^58^ suggests that this phytochemical richness contributes significantly to the biological/pharmacological activity potential of *Acacia* species.

### 3.3. Antimicrobial activity

Today, many emerging diseases (non-infectious and infectious) are treated using many plant-based medicines and has become an encouraging source of bioactives molecules and demonstrate a source of promising pharmacological agents. *Acacia* species is considered among the genera widespread uses as traditional as well as the modern medicine. In fact, several studies show that these species are rich in bioactive secondary metabolites such as polyphenols, flavonoids, terpenes, alkaloids ^59, 60^. On the other hand, many other important biological properties have been reported in *Acacia* species to treat a wide range of diseases such cancer, inflammation, diabetes, antioxidants, viral infections, and in the liver protection ^58^.

Antimicrobial activity of 100 mg/ml methanolic extracts from *A. gerrardii* were active against all tested bacterial and fungal species to varying degrees and were concentration dependent (Table 3, Figure 4). At a dose of 10 µL/disc (1mg extract), the mean growth inhibition zone (mGIZ) of the methanolic extract ranged from 6 to 9.33 mm for bacterial strains and about 6 mm for all tested yeast strains. The highest antibacterial activity was recorded against *K. pneumoniae* (9.33 ± 057 mm). The highest activity (highest mGIZ) was obtained at the highest dose, 30 µL/disc (3 mg) for all tested microorganisms. At 10 µL/disc, mGIZ ranged from 12.66 ± 0.57 mm (*A. baumannii* and *S. hominis*) to 15.33 ± 0.57 mm (*K. pneumoniae*) for the methanolic extract from Hail region. By the Duncan multiple range test, the antimicrobial activity is dependent on the concentration used (*p* < 0.05). The methanolic extract was also slightly active against yeast strains, and mGIZ ranged from 11.66 ± 0.57 mm (*C. guillermondii* ATCC 6260) to 14.66 ± 0.57 mm (*C. albicans* ATCC 20402).

**Figure 4.**
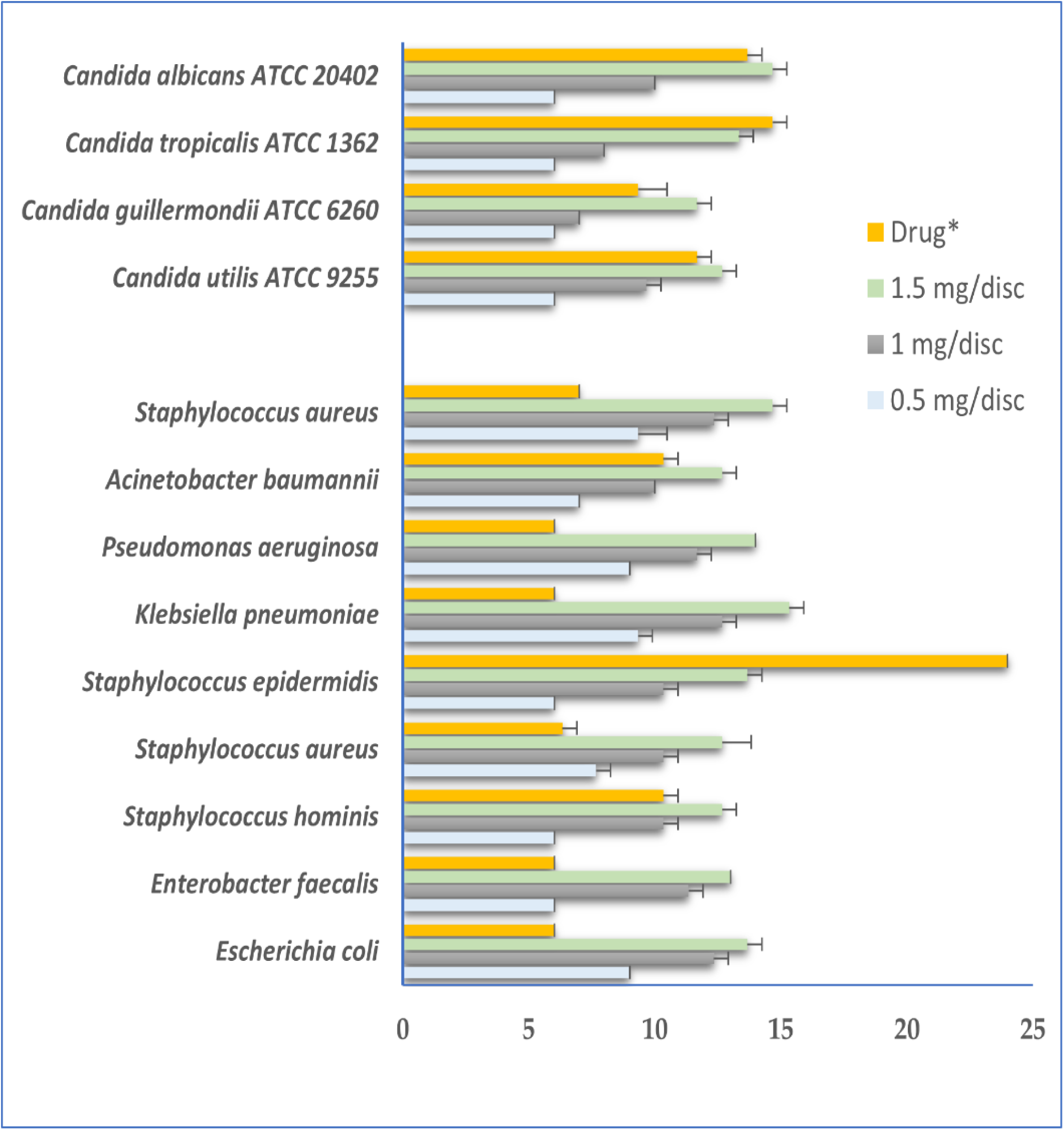
Mean diameters of bacterial and fungal growth inhibition zones (mGIZ±mm) obtained with different concentrations of methanolic extract as compared to standard drugs *: Ampicillin for bacteria and amphotericin B for *Candida* strains.

**Table 3.** Growth inhibition zone values expressed in mm of *A. gerrardii* methanolic extract tested against bacterial and yeast strains using disc diffusion assay.

| Code | Bacterial strain | Mean growth inhibition zone (mm) |  |  | Ampicillin |
| --- | --- | --- | --- | --- | --- |
|  |  | 10 µL/disc | 20 µL/disc | 30 µL/disc |  |
| M<sub>6</sub> | *E. coli* | 9 ± 0<sup>a,C</sup> | 12.33 ± 0.57<sup>a,b</sup> | 13.66 ± 0.57<sup>b,c,A</sup> | 6 ± 0<sup>d</sup> |
| M<sub>7</sub> | *E. faecalis* | 6 ± 0<sup>c,C</sup> | 11.33 ± 0.57<sup>b,c</sup> | 13 ± 0<sup>c,d,A</sup> | 6 ± 0<sup>d</sup> |
| M<sub>11</sub> | *S. hominis* | 6 ± 0<sup>c,C</sup> | 10.33 ± 0.57<sup>c,d</sup> | 12.66 ± 0.57<sup>d,A</sup> | 10.33 ± 0.57<sup>b</sup> |
| M<sub>12</sub> | *S. aureus* | 7.66 ± 0.57<sup>b,C</sup> | 10.33 ± 0.57<sup>c,d</sup> | 12.66 ± 1.15<sup>d,A</sup> | 6.33 ± 0.57<sup>d</sup> |
| M<sub>13</sub> | *S. epidermidis* | 6 ± 0<sup>c,C</sup> | 10.33 ±<br>0.57<sup>c,d,B</sup> | 13.66 ± 0.57<sup>b,c,A</sup> | 24 ± 0<sup>a</sup> |
| M<sub>14</sub> | *K. pneumoniae* | 9.33 ± 0.57<sup>a,C</sup> | 12.66 ± 0.57<sup>a,B</sup> | 15.33 ± 0.57<sup>a,A</sup> | 6 ± 0<sup>d</sup> |
| M<sub>16</sub> | *P. aeruginosa* | 9 ± 0<sup>a,C</sup> | 11.66 ±<br>0.57<sup>a,b,B</sup> | 14 ± 0<sup>b,c,A</sup> | 6 ± 0<sup>d</sup> |
| M<sub>17</sub> | *A. baumannii* | 7 ± 0<sup>b,C</sup> | 10 ± 0<sup>d,B</sup> | 12.66 ± 0.57<sup>d,A</sup> | 10.33 ± 0.57<sup>b</sup> |
| MRSA-217 | *S. aureus* | 9.33 ± 1.15<sup>a,C</sup> | 12.33 ± 0.57<sup>a,b</sup> | 14.66 ± 0.57<sup>a,b,A</sup> | 7 ± 0<sup>c</sup> |
| Code | Yeasts | 10 µL/disc | 20 µL/disc | 30 µL/disc | Amphotericin B |
| A<sub>1</sub> | *C. utilis* ATCC 9255 | 6 ± 0<sup>a,C</sup> | 9.66 ± 0.57<sup>a,B</sup> | 12.66 ± 0.57<sup>b,c,A</sup> | 11.66 ± 0.57<sup>c</sup> |
| A<sub>4</sub> | *C. guilliermondii* ATCC 6260 | 6 ± 0<sup>a,C</sup> | 7 ± 0<sup>c,B</sup> | 11.66 ± 0.57<sup>c,A</sup> | 9.33 ± 1.15<sup>b</sup> |
| A<sub>8</sub> | *C. tropicalis* ATCC 1362 | 6 ± 0<sup>a,C</sup> | 8 ± 0<sup>b,B</sup> | 13.33 ± 0.57<sup>b,A</sup> | 14.66 ± 0.57<sup>a</sup> |
| A<sub>15</sub> | *C. albicans* ATCC 20402 | 6 ± 0<sup>a,C</sup> | 10 ± 0<sup>a,B</sup> | 14.66 ± 0.57<sup>a,A</sup> | 13.66 ± 0.57<sup>a</sup> |

The lower MICs values ranged from 1.56 to 3.125 mg/mL against bacterial strains (Table 4). Using the scheme proposed by <u>Gatsing, Tchakoute, Ngamga, Kuiate, Tamokou,</u> NJI, Tchouanguep and Fodouop ^26^ and La, Bahi, Dje, Loukou and Guede-Guina ^61^, the tested extract exhibited bacteriostatic activity against almost all bacterial strains (MBC/MIC ratio > 4), while, the extracts exhibited bactericidal activity (MBC/MIC ratio ≤ 4) against *P. aeruginosa* (M16). Interestingly, methanolic extract from Hail region showed fungicidal activity toward all used *Candida* species with MFC/MIC ratio equal to 4.

**Table 4.** MICs, MBCs/MFCs expressed in mg/ml and MBCs/MIC, MFC/MIC ratio of *A. gerrardii* methanolic extract tested against bacterial and yeast strains (H: Hail)

| Code | Bacterial Strain | Microdilution assay H |  |  |
| --- | --- | --- | --- | --- |
|  |  | MIC | MBC | MBC/MIC ratio |
| M<sub>6</sub> | *E. coli* | 3.125 | 25 | 8; Bacteriostatic |
| M<sub>7</sub> | *E. faecalis* | 3.125 | 25 | 8; Bacteriostatic |
| M<sub>11</sub> | *S. hominis* | 1.56 | 12.5 | 8; Bacteriostatic |
| M<sub>12</sub> | *S. aureus* | 1.56 | 12.5 | 8; Bacteriostatic |
| M<sub>13</sub> | *S. epidermidis* | 3.125 | 25 | 8; Bacteriostatic |
| M<sub>14</sub> | *K. pneumoniae* | 3.125 | 25 | 8; Bacteriostatic |
| M<sub>16</sub> | *P. aeruginosa* | 3.125 | 12.5 | 4; Bactericidal |
| M<sub>17</sub> | *A. baumannii* | 3.125 | 25 | 8; Bacteriostatic |
| MRSA-217 | *S. aureus* | 3.125 | 50 | 16; Bacteriostatic |
| Code | Yeasts | MIC | MFC | MFC/MIC ratio |
| A<sub>1</sub> | *C. utilis* ATCC 9255 | 6.25 | 25 | 4; Fungicidal |
| A<sub>4</sub> | *C. guilliermondii*<br>ATCC 6260 | 12.5 | 50 | 4; Fungicidal |
| A<sub>8</sub> | *C. tropicalis* ATCC<br>1362 | 12.5 | 50 | 4; Fungicidal |
| A<sub>15</sub> | *C. albicans* ATCC<br>20402 | 12.5 | 50 | 4; Fungicidal |

There are some studies analyzing the antimicrobial activity of honey produced from *A. gerrardii* ^62, 63^. However, there are no reports on the antimicrobial activity of extracts from the tree itself. However, there are reports of the antimicrobial activities of different *Acacia* species such as *A. nilotica* ^40, 41^, *A. ataxacantha* ^51^, *A. plicosepalus* ^52^, *A. farnesiana* ^64^ and *A. rigidula* ^57^. These studies suggest that phentolamine, betulinic acid, betulinic acid-3-*trans*-caffeate, loranthin, quercetin, methyl gallate, diterpenes and tannins, which stand out as the main components, may be the phytochemicals responsible for antimicrobial activity.

### 3.4. Antioxidant activity

Free radicals produced by oxidative processes or by intermediates occurring during metabolic reactions are the cause of many chronic metabolic diseases. Plants protect themselves by producing molecules to prevent free radical damage that are also useful to human health eliminate the damage caused by free radicals to the body ^65^. Many tests have been developed to measure the activity level of these antioxidant compounds. Some tests measure the antioxidant’s ability to inhibit lipid oxidation, while others document its free radical scavenging, metal chelating or reducing power efficiencies. In order to accurately evaluate the antioxidant activity of the target substance, several of these methods must be used in combination ^66^. The antioxidant activities of the methanol extract obtained from *A. gerrardii* leaves were evaluated using DPPH radical scavenging and FRAP tests.

The DPPH radical scavenging efficiency of the extract was determined as 0.28 ± 0.0045 mg/mL (IC_50_). Although this value is not as high as the radical scavenging activity of BHT used as a positive control, the extract exhibits significant radical scavenging activity. A similar situation applies to the FRAP test. In this system, the reducing power of the methanol extract was determined as 63 63 ± 1.12 mg/mL (IC_50_), while the reducing capacity of ascorbic acid used as a positive control agent was determined as 55.3 ± 1.21 mg/mL. There are some studies in the literature that measure the effect of changes in environmental stress factors on the antioxidant defense mechanism of *A. gerrardii* ^67, 68^. In addition, in the literature, there are some reports investigating the antioxidant activities of *A. ataxacantha*, which has high betulinic acid and betulinic acid-3-*trans*-caffeate contents ^51^, *A. plicosepalus*, which has high loranthin and quercetin content ^52^, *A. arabica*, which has high quercetine 3-*O*-(40-*O*-acetyl)-rhamnopyranoside content ^53^, *A. cyanophylla*, which has high naringenin content ^54^, *A. crassicarpa*, which has high quercetin,5,7,20,50-tetrahydroxyflavone content ^55^, *A. saligna*, which has high myricetin- 3-*O*-α-L-rhamnoside and quercetin-3-*O*-α-L-rhamnoside contents ^56^, and *A. pennatula*, which has high 4-nitro-ophenylenediamine content ^69^. However, there are no studies investigating the antioxidant activity of extracts or individual compounds obtained from *A. gerrardii* on other organism systems or *in vitro*.

Although it is not possible to compare the antioxidant activity of the plant with literature data, it is possible to get an idea about the contributions of the compounds found in high amounts in the methanol extract to antioxidant activity. In a study by <u>Sousa,</u> Gomes, Viana, Silva, Barata, Sartoratto, Lustosa and Vieira ^70^, the chemical composition and antioxidant activities of extracts obtained from different parts of *Dipteryx punctata* were analyzed and it was reported that residue extracts containing high amounts of 4-*O*- methylmannose exhibited the highest DPPH radical scavenging activity. Another study emphasized that *Portulaca oleracea* is a notable antioxidant due to its high α-linolenic acid content and may be an important element of the human diet ^71^. On the other hand, *Achillea filipendulina*, which contains high amounts of 13-docosenamide, *(Z)* as the main component, scavenged DPPH free radical by 92.98% ^72^.

### 3.5. Computational pharmacokinetic analyses

The lipophilicity, pharmacokinetics and bioavailability properties of the compounds identified in *A. gerrardii* are shown in Table 5. Most of the A. gerrardii identified phytochemicals meet the Lipinski rule for therapeutic properties. Lipophilicity (lipo), polarity (pola), molecular size (size), insolubility (insolu), insaturation (insatu) and flexibility (flex) of A. gerrardii phytochemicals follow the Lipinski rule, are suitable for oral bioavailability,and, hence, confirmed the potential biological activities of the extract (Figure 5). Gastro-intestinal (GI) absorption and the blood-brain-barrier (BBB) permeation of *A. gerrardii* compounds have also been assessed. GI absorption and BBB permeation were evaluated by calculating n-octanol/water partition coefficient (log P) and the polar surface area (PSA) and applying them using the boiled egg-model (Figure 6). Nine compounds fell into the BBB permeable area of the boiled egg model. Most of the remainder had properties suggesting good gastrointestinal permeability. P-gp is a ATP binding cassette transporter removing xenobiotics and toxins from the cell. Most of the *A. gerardii* compounds are not substrate of P-glycoprotein (P-gp) and would persist in the cell. Cytochromes P450 (CYPs) have a key role in the metabolism and excretion of drugs and xenobiotics. The possible inhibition of several CYPs (1A2, 2C19, 2C9, 2D6 and 3A4) was also assessed by computational study. Our findings showed that most of the compounds did not inhibit the assessed CYPs, which indicate no disruptions of metabolism and excretion. Only CYP2D6 and CYP3A4 were inhibited by 3 compounds and CYP2C19 was inhibited only by compounds 13 and 43. Levels of log Kp, an index of skin permeation, ranged between –3.23 and –13.76 kcal/mol supporting moderate to low permeation. The synthetic accessibility of *A. gerrardii* phytochemicals varied between 1.77 and 5.61.

**Figure 5.**
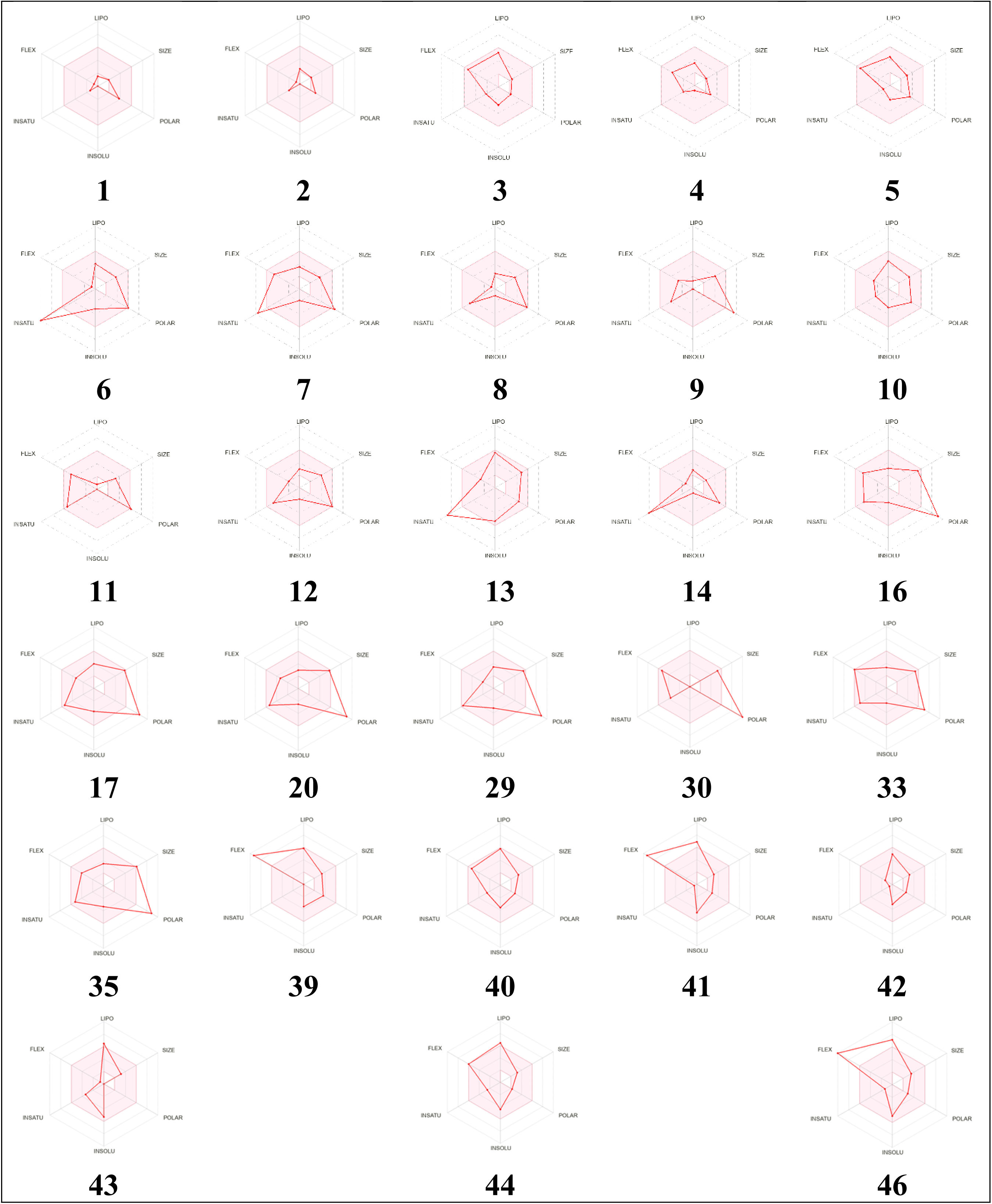
Bioavailability hexagons of the identified compounds of *A. gerrardii* based on their physicochemical properties; lipophilicity (Lipo), molecular size (Size), polarity (Pola), insolubility (Insolu), insaturation (Insatu) and flexibility (Flex)

**Figure 6.**
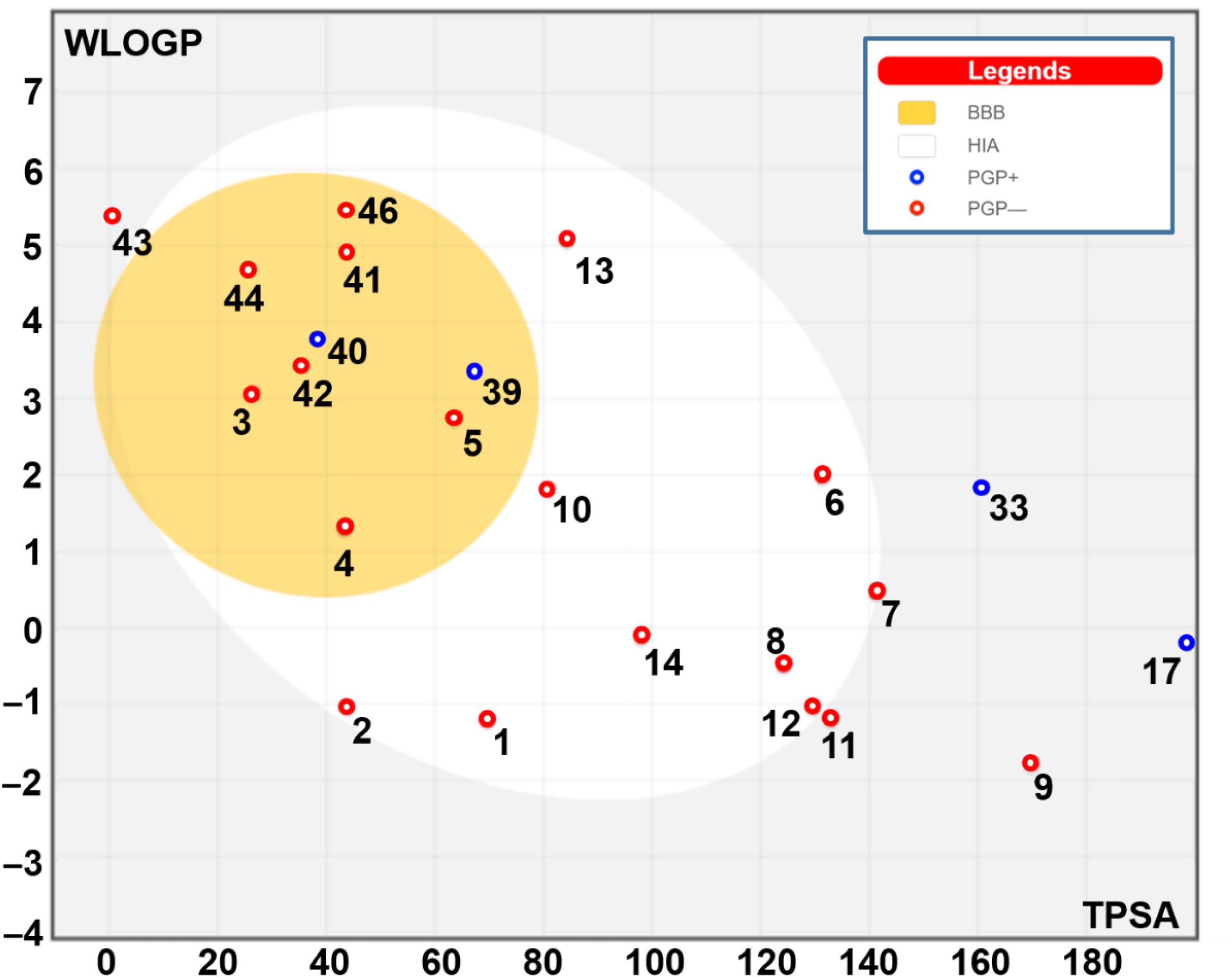
Boiled-egg model of the identified compounds of *A. gerrardii* (1-17) based on their GI absorption, BBB permeation and interaction with P-gp properties Note that some compounds are out of range.

**Table 5.** Lipophilicity, pharmacokinetics, druglikeness and medicinal chemistry of the identified compounds in *A. gerrardii* based on their ADMET (for absorption, distribution, metabolism, excretion and toxicity) properties.

| Entry | 1 | 2 | 3 | 4 | 5 | 6 | 7 | 8 | 9 | 10 | 11 | 12 | 13 | 14 | 16 |
| --- | --- | --- | --- | --- | --- | --- | --- | --- | --- | --- | --- | --- | --- | --- | --- |
|  | **Lipophilicity & Physicochemical properties** |  |  |  |  |  |  |  |  |  |  |  |  |  |  |
| TPSA | 69.56 | 43.70 | 26.30 | 43.37 | 63.60 | 131.36 | 141.36 | 124.29 | 169.86 | 80.67 | 132.88 | 129.59 | 84.18 | 97.99 | 215.58 |
| Log *P*<sub>o/w</sub> (iLOGP) | 0.81 | 1.45 | 3.21 | 2.16 | 2.34 | 1.63 | 0.51 | 0.77 | -0.21 | 1.70 | 0.17 | 1.65 | 2.98 | 0.58 | 0.37 |
| Consensus Log *P*<sub>o/w</sub> | -1.34 | -0.21 | 3.15 | 1.32 | 2.43 | 1.23 | 0.38 | -0.30 | -2.12 | 1.92 | -1.25 | -0.17 | 4.19 | -0.12 | -0.45 |
| Log *S* (ESOL) solubility | 1.10 | 0.08 | -2.77 | -0.80 | -2.23 | -3.16 | -1.82 | -1.06 | 0.08 | -2.94 | 1.32 | -1.82 | -5.30 | -0.84 | -2.33 |
|  | **Pharmacokinetics** |  |  |  |  |  |  |  |  |  |  |  |  |  |  |
| GI absorption | High | High | High | High | High | High | High | High | Low | High | Low | High | Yes | High | Low |
| BBB permeant | No | No | Yes | Yes | Yes | No | No | No | No | No | No | No | No | No | No |
| P-gp substrate | No | No | No | No | No | No | No | No | No | No | No | No | No | No | No |
| CYP1A2 | No | No | No | No | No | Yes | No | No | No | No | No | No | Yes | No | No |
| CYP2C19 | No | No | No | No | No | No | No | No | No | No | No | No | Yes | No | No |
| CYP2C9 | No | No | No | No | No | No | No | No | No | No | No | No | Yes | No | No |
| CYP2D6 | No | No | No | No | No | Yes | No | No | No | No | No | No | No | No | No |
| CYP3A4 | No | No | No | No | No | Yes | No | No | No | No | No | No | No | No | No |
| Log *K*<sub>p</sub> (skin permeation) | -9.18 | -8.16 | -4.82 | -6.84 | -6.18 | -7.05 | -7.67 | -8.94 | -10.60 | -6.55 | -10.86 | -8.46 | -5.68 | -7.82 | -9.10 |
|  | **Drug likeness & Medicinal chemistry** |  |  |  |  |  |  |  |  |  |  |  |  |  |  |
| Lipinski | Yes | Yes | Yes | Yes | Yes | Yes | Yes | Yes | Yes | Yes | Yes | Yes | Yes | Yes | No |
| Ghose | No | No | Yes | No | Yes | Yes | Yes | No | No | Yes | No | No | Yes | Yes | Yes |
| Veber | Yes | Yes | Yes | Yes | Yes | Yes | No | Yes | No | Yes | Yes | Yes | Yes | Yes | No |
| Ergan | Yes | Yes | Yes | Yes | Yes | Yes | No | Yes | No | Yes | No | Yes | Yes | Yes | No |
| Muegge | No | No | No | No | Yes | Yes | Yes | Yes | No | Yes | No | Yes | Yes | No | No |
| Bioavailability score | 0.55 | 0.55 | 0.55 | 0.55 | 0.85 | 0.55 | 0.56 | 0.56 | 0.11 | 0.55 | 0.55 | 0.55 | 0.55 | 0.56 | 0.11 |
| Synthetic accessibility | 2.31 | 3.47 | 2.52 | 1.77 | 3.48 | 3.23 | 3.15 | 4.50 | 4.18 | 4.70 | 2.47 | 4.68 | 2.32 | 1.95 | 5.16 |

TPSA: Topological polar surface area, GI: Gastro-intestinal, BBB: Blood-brain-barrier, P-gp: P-glycoprotein, CYP: Cytochrome P450

| Entry | 17 | 20 | 29 | 30 | 33 | 35 | 39 | 40 | 41 | 42 | 43 | 44 | 46 |
| --- | --- | --- | --- | --- | --- | --- | --- | --- | --- | --- | --- | --- | --- |
|  | **Lipophilicity & Physicochemical properties** |  |  |  |  |  |  |  |  |  |  |  |  |
| TPSA | 199.51 | 212.67 | 210.51 | 248.20 | 160.48 | 208.74 | 66.48 | 38.83 | 43.09 | 35.53 | 0.00 | 26.30 | 43.09 |
| Log Po/w (iLOGP) | 1.83 | 0.16 | 1.71 | 0.08 | 1.98 | 2.54 | 3.52 | 3.52 | 3.87 | 3.23 | 3.59 | 4.21 | 4.10 |
| Consensus Log Po/w | 0.12 | -0.67 | -0.07 | -3.69 | 1.25 | 0.07 | 3.64 | 3.64 | 4.83 | 3.15 | 5.38 | 4.64 | 5.29 |
| Log S (ESOL) solubility | -3.73 | -2.53 | -3.20 | 2.04 | -2.47 | -3.34 | -3.59 | -3.59 | -4.61 | -3.21 | -5.28 | -4.46 | -5.00 |
|  | **Pharmacokinetics** |  |  |  |  |  |  |  |  |  |  |  |  |
| GI absorption | Low | low | Low | Low | Low | Low | High | High | High | High | Low | High | High Yes |
| BBB permeant | No | No | No | No | No | No | Yes | Yes | Yes | Yes | No | Yes | Yes |
| P-gp substrate | Yes | No | No | No | Yes | Yes | Yes | Yes | No | No | No | No | No |
| CYP1A2 | No | No | No | No | No | No | No | No | Yes | No | No | Yes | Yes |
| CYP2C19 | No | No | No | No | No | No | No | No | No | No | Yes | No | No |
| CYP2C9 | No | No | No | No | No | No | No | No | No | No | Yes | Yes | Yes |
| CYP2D6 | No | No | No | No | No | No | Yes | Yes | No | No | No | No | No |
| CYP3A4 | Yes | No | No | No | No | Yes | No | No | No | No | No | No | No |
| Log Kp (skin permeation) | -8.20 | -9.48 | -8.77 | -13.76 | -8.84 | -8.93 | -4.62 | -4.62 | -3.23 | -5.68 | -3.69 | -3.72 | -3.05 |
|  | **Drug likeness & Medicinal chemistry** |  |  |  |  |  |  |  |  |  |  |  |  |
| Lipinski | No | No | No | No | Yes | No | Yes | Yes | Yes | Yes | Yes | Yes | Yes |
| Ghose | Yes | No | No | No | Yes | No | Yes | Yes | Yes | Yes | Yes | Yes | Yes |
| Veber | No | No | No | No | No | No | No | No | No | Yes | Yes | Yes | No |
| Ergan | No | No | No | No | No | No | Yes | Yes | Yes | Yes | Yes | Yes | Yes |
| Muegge | No | No | No | No | No | No | Yes | Yes | No | Yes | No | No | No |
| Bioavailability Score | 0.17 | 0.17 | 0.17 | 0.17 | 0.11 | 0.17 | 0.55 | 0.55 | 0.55 | 0.55 | 0.55 | 0.55 | 0.55 |
| Synthetic accessibility | 5.30 | 4.95 | 5.32 | 5.04 | 4.42 | 5.61 | 3.19 | 3.19 | 2.12 | 4.23 | 3.19 | 3.15 | 2.97 |

TPSA: Topological polar surface area, GI: Gastro-intestinal, BBB: Blood-brain-barrier, P-gp: P-glycoprotein, CYP: Cytochrome P450

The identified compounds of *A. gerrardii* showed different affinities to TyrRS from methicillin resistant *S. aureus* (1JIJ) and aspartic proteinase from the pathogenic yeast, *C. albicans* (2QZW), receptors (Table 6). All of them had negative binding affinities that would support their biological potentialities to inhibit growth of these microorganisms. The binding scores varied between −4.7 and −10.4 kcal/mol for 1JIJ, and between −4.3 and −9.9 kcal/mol for 2QZW. Such variations have been reported to be related to the 3D chemical structure of both ligand and receptors (Rahmouni et al., 2022; Ben Saad et al., 2023).The best binding scores were predicted for compounds 20 and 29, while complexed with 1JIJ with −10.0 and −10.4 kcal/mol (Table 7). Compound 29 was predicted to establish excellent molecular interactions with the 1JIJ that included 9 conventional H-bonds together with a network of carbon H-bonds, electrostatic, alkyl and Pi-alkyl bonds, which concerned several key residues. In fact, it involved eleven different residues Cys37, Asp40, Lys84, Gly193, Gly38, Asp195, His50, Pro53, Phe54, Ala39, and Tyr170 (Figures 7 and 8). This compound, similarly to several others, was deeply embedded in the pocket region of 1JIJ and showed a distance of 1.965 Å only.

**Figure 7.**
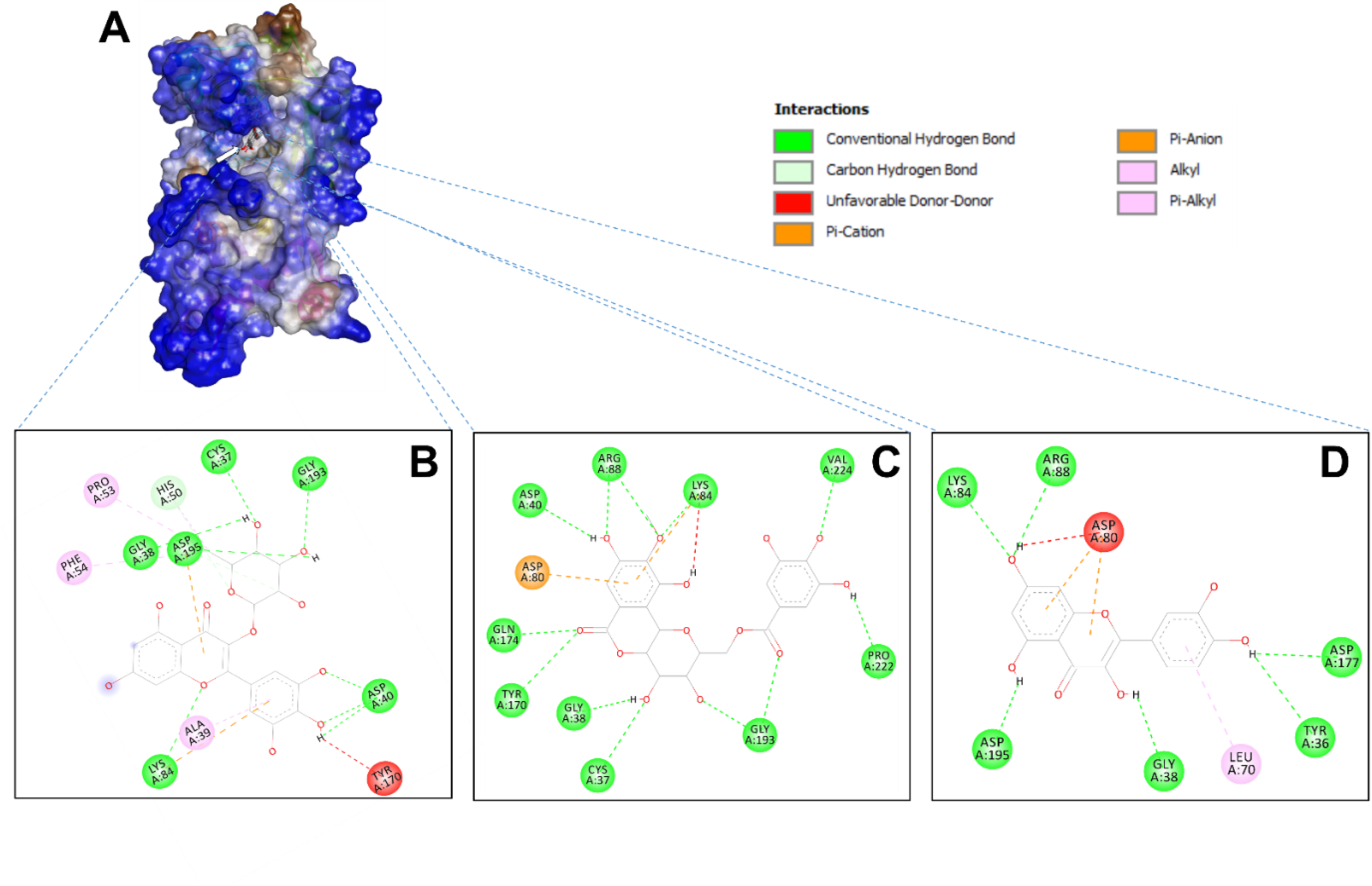
3D illustration of the hydrophobic complex 1JIJ-ligand (A) and the resulting 2D diagram of interactions of compounds 29 (B), 20 (C) and 6 (D) of *A. gerrardii* that possessed the best predicted binding affinities (–10.4, –10.0, and –9.9, respectively)

**Figure 8.**
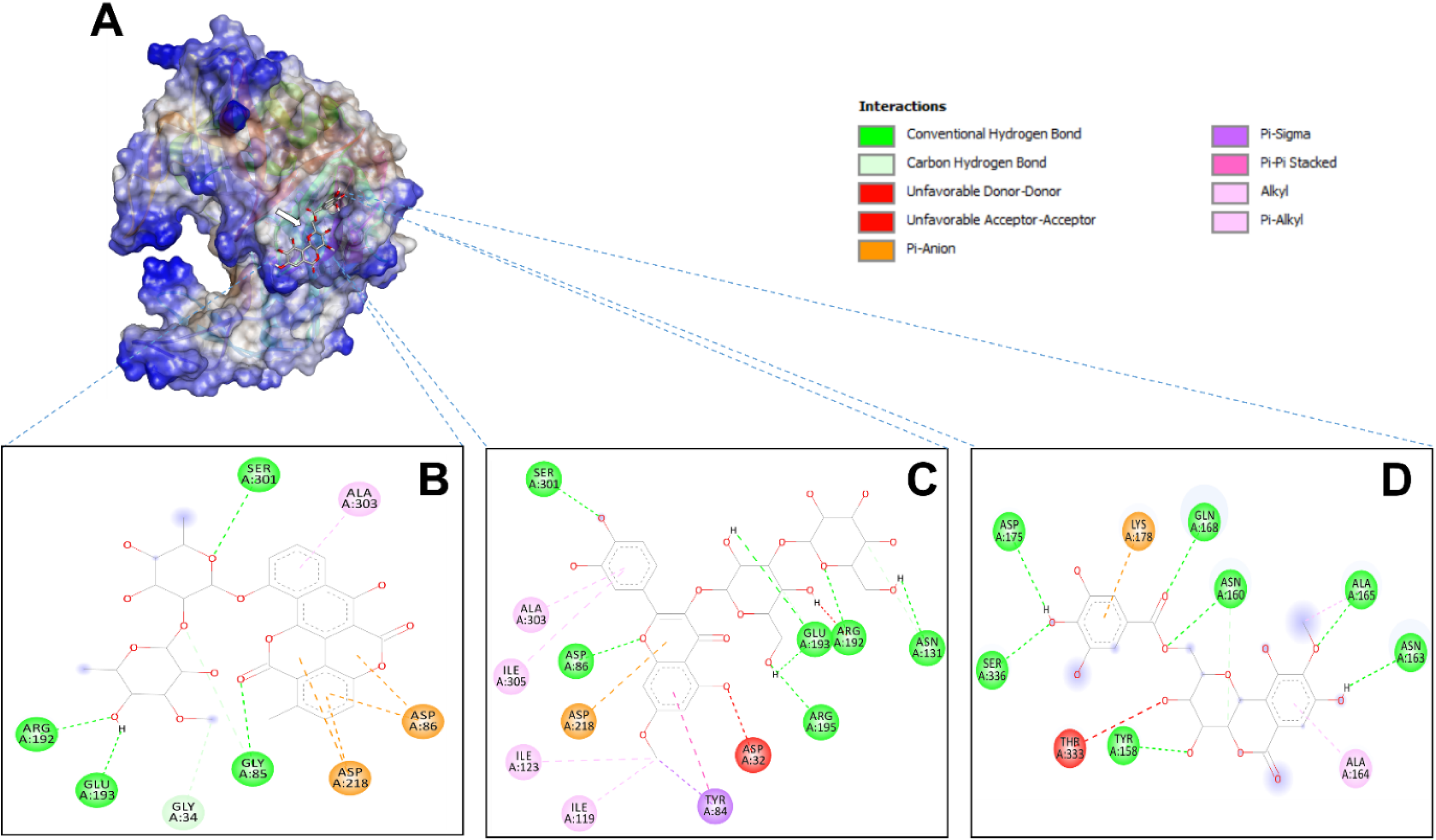
3D illustration of the hydrophobic complex 2QZW-ligand (A) and the resulting 2D diagram of interactions of compounds 21 (B), 23 (C) and 20 (D) of *A. gerrardii* that possessed the best predicted binding affinities (–9.9, –9.4i and –9.1, respectively)

**Table 6.**
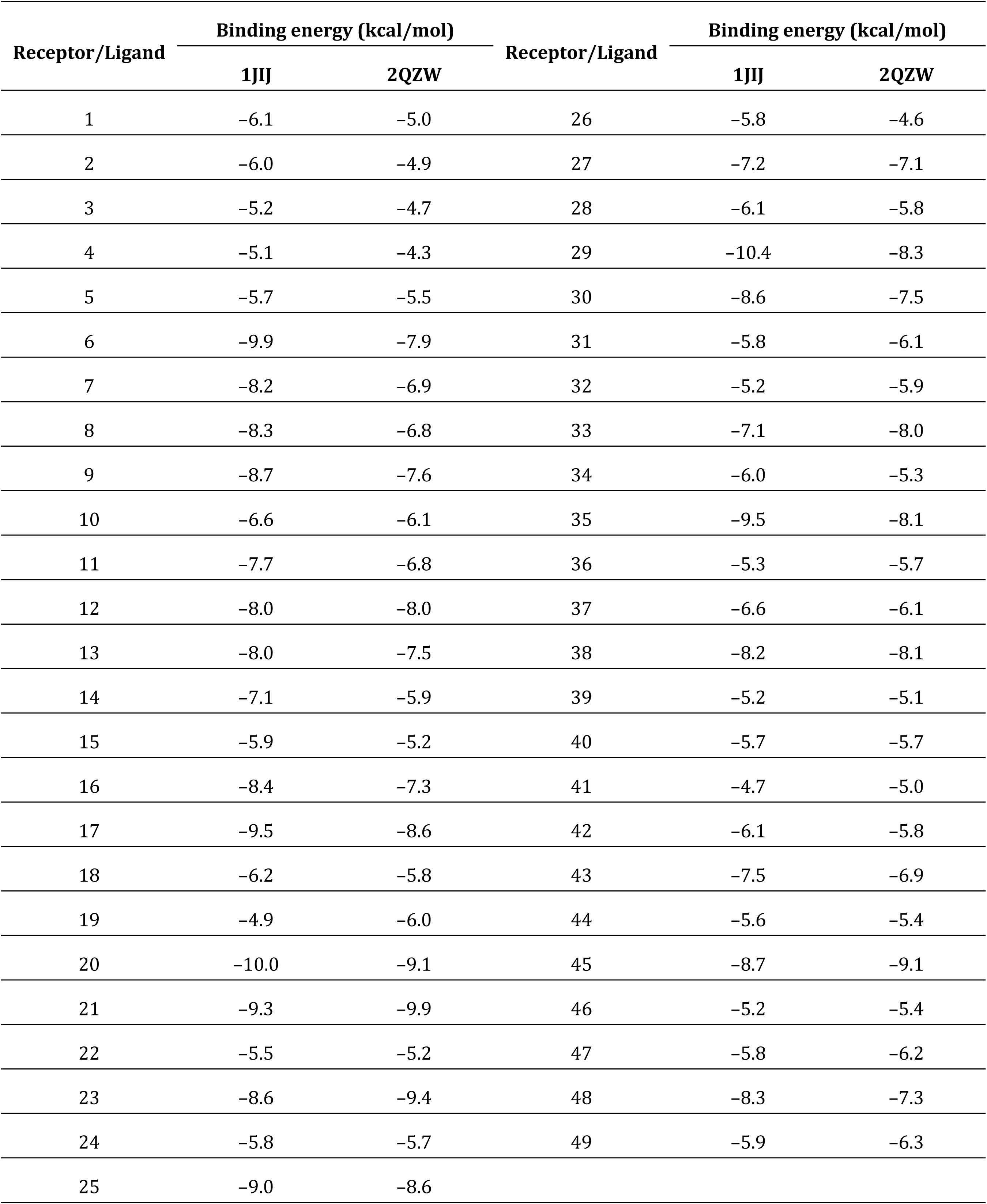
Binding energy of the identified compounds of *A. gerrardii* to the two targeted receptors: 1JIJ and 2QZW for TyrRS from *S. aureus* and aspartic proteinase from *C. albicans*, respectively.

**Table 7.**
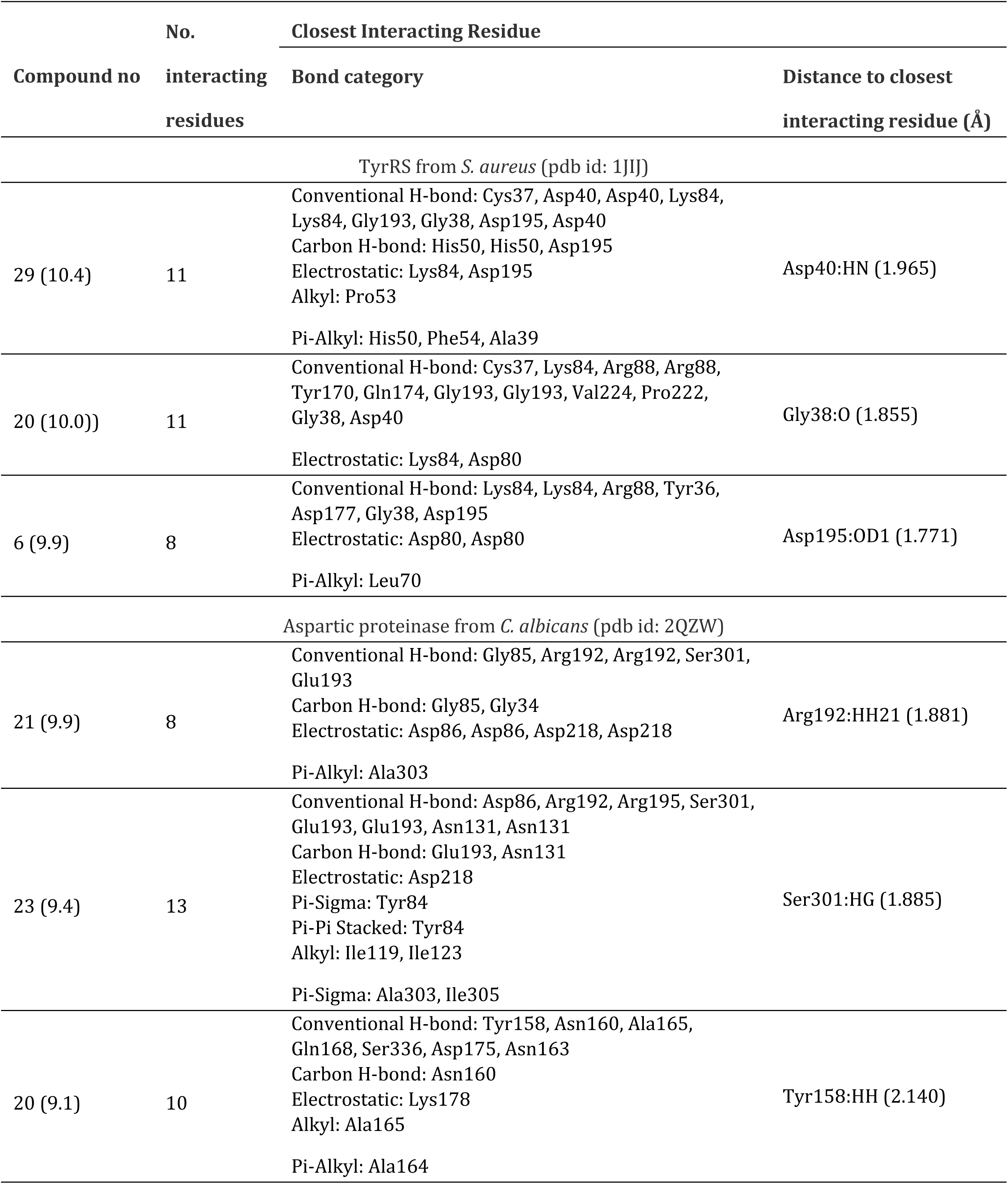
Interactions, bond category and closest interacting residues for the best *A. gerrardii* compounds with the targeted receptors: 1JIJ and 2QZW for TyrRS from *S. aureus* and aspartic proteinase from *C. albicans*, respectively.

The assessment of lipophilicity, pharmacokinetic and bioavailability properties is commonly studied to avoid drug failure in the advanced stages of drug design and development ^30, 33, 34^. The *A. gerrardii* identified phytochemicals were predicted to meet the Lipinski rule and possessed druglikeness properties. They also possessed suitable for oral bioavailability and, hence, confirmed the potential biological activities of the extract. Gastro-intestinal (GI) absorption and the blood-brain-barrier (BBB) permeation of *A. gerrardii* compounds have also been assessed. GI absorption and BBB permeation were used for the mapping of the egg-model ^33^. Good oral bioavailability is commonly associated with significant biological effects for natural and synthesized compounds ^30, 73^. Furthermore, the majority of the *A. gerrardii* phytochemicals were predicted not being substrate of P-glycoprotein (P-gp), which supports their safe use and the absence of any possible toxicological outcome. Cytochromes P450 (CYPs) have a key role in the metabolism and excretion of drugs ^35, 74, 75^; Bédoui et al., 2023. Studying the possible inhibition of CYPs, the results revealed that the first several compounds inhibited none of the assessed CYPs, which indicate no metabolism and excretion disruptions. Moreover, CYP2D6 and CYP3A4 were inhibited by 3 compounds only and CYP2C19 was inhibited only by compounds 13 and 43. *A. gerrardii* phytochemicals are easy to synthesize as the synthetic accessibility varied between 1.77 and 5.61 ^30, 74, 75^ <u>Bédoui et al., 2023</u>.

*A. gerrardii* compounds had various affinities to TyrRS from *S. aureus* (1JIJ) and aspartic proteinase from *C. albicans* (2QZW) receptors (Table 6). Recent studies reported that variations in binding affinities are related to both chemical structure of the compounds and the structural geometry of the ligands ^30, 31, 33^. In this study, all the 49 compounds of *A. gerrardii* had negative binding affinities that would support their biological potentialities. The binding scores varied between −4.7 and −10.4 kcal/mol for 1JIJ, and between −4.3 and −9.9 kcal/mol for 2QZW. The best binding scores were predicted for compounds 20 and 29 while complexed with 1JIJ with −10.0 and −10.4 kcal/mol (Table 7). Compound 29 established good molecular interactions with the 1JIJ that included 9 conventional H-bonds associated with a network of electrostatic, alkyl and Pi-alkyl bonds that enhance the stability of the complex ^30, 31, 34^. These intermolecular interactions involved several key residues. In fact, it involved eleven different residues Cys37, Asp40, Lys84, Gly193, Gly38, Asp195, His50, Pro53, Phe54, Ala39 and Tyr170 (Figure 7). This compound, similarly to several others, was also tightly embedded in the pocket region of 1JIJ and showed a distance of 1.965 Å only. Tight embedding (<2.5 Å), as those of the current study, was commonly reported to be actively involved in several biological activities including antiinflammatory, antiproliferative and antimicrobial effects ^32, 74, 76^. Taken together, the binding affinities, the deep embedding and the established molecular interactions of *A. gerrardii* phytochemicals indicate that the antibacterial and antiviral effects are thermodynamically possible. Both effects had already been approved experimentally using in vitro analyses. Thus, supporting the promising health promotion and beneficial effects of natural-derived compounds, phytotherapy and medicinal plants ^33, 74, 76^.

The methanolic extract obtained from the leaves of *A. gerradii* exhibited antimicrobial activity toward the tested bacterial and fungal strains. It also showed high antioxidant activity using DPPH radical scavenging and FRAP tests. The pharmacokinetic properties, the binding affinities of the compounds to the target residues, disk diffusion, MIC/MBC tests and the results of *in silico* analysis supports the antimicrobial properties of natural products found in *A. gerradii*. GC-MS and HR-LCMS analysis to identify the natural products of the methanolic extract of *A. gerradii*. In addition to its therapeutic natural products, Acacia species are of great importance for the agricultural industry of the Hail region of Saudi Arabia due to their synbiotic relationships to revitalize soil nitrogen.

## Declaration of Competing Interest

The authors confirm that there are no known conflicts of interest.

## Acknowledgment

This research has been funded by Deputy for Research & Innovation, Ministry of Education through Initiative of Institutional Funding at University of Ha’il – Saudi Arabia through project number IFP 22-051”.

## CRediT authorship contribution statement

**Salem ELKAHOUI**: Conceptualization, Methodology, Data curation, Writing – original draft. **Ahmed Eisa Mahmoud Ghoniem**: Methodology, Resources. **Mejdi Snoussi**: Supervision, Methodology, Writing – review & editing. **Zohaier Barhoumi**: Resources, Data analysis (GC and LC/MS). **Riadh Badraoui**: Software, “in silico” analysis, Writing - review & editing.

## Notes

### Competing Interest Statement

The authors have declared no competing interest.

